# Liquid footprinting: A novel approach for DNA footprinting using short double-stranded cell-free DNA from plasma

**DOI:** 10.1101/2024.02.09.579588

**Authors:** Jan Müller, Christina Hartwig, Mirko Sonntag, Lisa Bitzer, Christopher Adelmann, Yevhen Vainshtein, Karolina Glanz, Sebastian O. Decker, Thorsten Brenner, Georg F. Weber, Arndt von Haeseler, Kai Sohn

**Affiliations:** Innovation Field In-vitro Diagnostics, Fraunhofer Institute for Interfacial Engineering and Biotechnology IGB, Stuttgart, Germany; Max Perutz Labs, Vienna Biocenter Campus (VBC), Vienna, Austria; University of Vienna, Max Perutz Labs, Department of Structural and Computational Biology, CIBIV, Vienna, Austria; Vienna BioCenter PhD Program, Doctoral School of the University of Vienna and Medical University of Vienna, Vienna, Austria; Institute for Interfacial Engineering and Plasma Technology (IGVP), University of Stuttgart, Stuttgart, Germany; Interfaculty Graduate School of Infection Biology and Microbiology (IGIM), Eberhard Karls University Tübingen, Tübingen, Germany; Heidelberg University, Medical Faculty Heidelberg, Department of Anesthesiology, Heidelberg, Germany; Department of Anesthesiology and Intensive Care Medicine, University Hospital Essen, University Duisburg-Essen, Essen, Germany; Department of Surgery, Friedrich-Alexander University (FAU) Erlangen-Nürnberg and Universitätsklinikum Erlangen, Erlangen, Germany; Comprehensive Cancer Center (CCC) Erlangen-EMN, Friedrich-Alexander University (FAU) Erlangen-Nürnberg and Universitätsklinikum Erlangen, Erlangen, Germany; Center of Integrative Bioinformatics Vienna (CIBIV), Max Perutz Labs, University of Vienna and Medical University of Vienna, Vienna BioCenter (VBC), Vienna, Austria; University of Vienna, Faculty of Computer Science Bioinformatics and Computational Biology, Vienna, Austria

## Abstract

The diagnostic potential of short double-stranded cell-free DNA (cfDNA) in blood plasma has not been recognized, yet. Here, we present a method for enrichment of double-stranded cfDNA with an average length of about 40 base pairs from cfDNA for high-throughput DNA sequencing. This class of cfDNA is enriched at gene promoters and binding sites of transcription factors or structural DNA-binding proteins, so that a genome-wide DNA footprint is directly captured from liquid biopsies. In short double-stranded cfDNA from healthy individuals, we found significant enrichment of 203 transcription factor motifs. Additionally, short double-stranded cfDNA signals at specific genomic regions correlate negatively with DNA-methylation, positively with H3K4me3 histone modifications and gene transcription. When comparing short double-stranded cfDNA from patient samples of pancreatic ductal adenocarcinoma with colorectal carcinoma or septic with post-operative controls, we identified 731 and 1,107 differentially enriched loci, respectively. Using these differentially enriched loci, the disease types can be clearly distinguished by principal component analysis demonstrating the diagnostic potential of short double-stranded cfDNA signals as a new class of biomarkers for liquid biopsies.

## Introduction

Liquid biopsies are based on various types of analytes, including circulating extracellular nucleic acids like cell-free DNA (cfDNA), extracellular vesicles, or circulating tumor cells, for example^1,2^. Physiologically, cfDNA is to large extend released from the hematopoietic system by apoptosis, necrosis, or active secretion from almost all cell types and tissues into the bloodstream^3,4^. In addition to release from normal physiological turnover, cancer cells or microbial pathogens are also known to release their DNA into bloodstream circulation^3^. Released genomic DNA is then degraded by DNA-digesting enzymes (nucleases), producing fragments mainly 147 to 167 base pairs (bp) in size, corresponding to a single nucleosome^4,5^. By high throughput sequencing of cfDNA fragments, nucleosome positioning can be inferred at base pair resolution^6^. The exact positions of nucleosomes and chromatin structure play a key role in regulating gene expression by providing access to DNA for the transcription machinery. Open chromatine structures depleted of nucleosomes make DNA more accessible for key regulators of transcription, including transcription factors, enhancers, or repressors. However, the routine and efficient measurement of genome-wide protection through regulatory DNA-binding proteins (DBP) is not yet established. Recently, a minor fraction of double-stranded cfDNA that is significantly shorter than normal cfDNA was found, ranging from 35 to 80 nucleotides using ultra-deep sequencing of total cfDNA^6,7^. It has been proposed that this short cfDNA might be protected by DNA-binding factors and therefore could represent direct transcription factor binding.

Here, we established an enrichment approach for short double-stranded cfDNA fragments (20-60 bp) from blood plasma (‘Liquid Footprinting’). Corresponding short double-stranded cfDNA fragments (further referred to as ‘footprint DNA’) accumulate at open chromatin as well as gene regulatory elements. Differential enrichment of footprint DNA at genomic loci facilitated discrimination of colorectal and pancreatic cancer patient samples as well as between septic patients and clinical controls showing potential for diagnostic applications.

## Results

### Enrichment of cfDNA footprints at regulatory genomic regions

Enrichment of short double-stranded cfDNA comprises extraction of total cfDNA from blood plasma by magnetic beads followed by double-stranded DNA-specific library preparation (**Supplementary Fig. S1**). To select short double-stranded cfDNA fragments of up to 60 bp, two size selection steps were performed using a preparative gel electrophoresis device (**Supplementary Fig. S2**). After high-throughput sequencing of size-selected libraries, reads were quality-checked to ensure a size range between 20 bp and 60 bp (**Supplementary Fig. S3**). The protocol revealed sequencing reads with a mean read length of 37.9 bp (SD = 6.6 bp, n = 2·10^7^ reads uniformly sampled from the total reads of 20 individuals: four healthy individuals, four patients with pancreatic ductal adenocarcinoma (PDAC), four patients with colorectal carcinoma (CRC), four patients with sepsis, and four non-septic post-operative clinical control patients (Post-OP); **Supplementary Fig. S4; Supplementary Table 1**). Footprint DNA reads revealed an elevated average GC content of 57.8 % (SD = 1.9 %, **Supplementary Fig. S5**) in contrast to 40.9 % for average human genomic DNA^8^. We mapped footprint DNA to the human genome and compared it with the mapping of regular cfDNA of four additional healthy individuals. Coverage profiles of footprint DNA and regular cfDNA differed considerably as footprint DNA tends to accumulate either in single narrow peaks or in clusters of narrow peaks (clustered peaks), which are frequently located at transcriptional start sites (TSS) or ChIP-Seq validated transcription factor binding sites (**Fig. 1a**). Nucleosome-free regions (NFRs) were analyzed in comparison to clustered peaks, since the absence of regular cfDNA can be an indicator of the presence of other DBPs. Assignment of clustered peaks from footprint DNA and NFRs to annotated functional elements of the genome shows that an approximately four- and a six-fold higher proportion of clustered peaks are found in promoters (>1 Kb upstream of the TSS) and 5’ UTR of genes, respectively. Moreover, the proportion of clustered peaks assigned to exons is about four times higher than the proportion of NFRs (**Fig. 1b**). A more detailed examination of the genomic coverage for protein-coding genes reveals an opposite profile between footprint DNA and regular cfDNA. While footprint DNA is strongly enriched at TSS, regular cfDNA is substantially depleted at these sites. In addition, footprint DNA possesses a reciprocal pattern oscillation compared to regular cfDNA, with footprint DNA exhibiting an inverted pattern 1 Kb downstream of the TSS in regular cfDNA (insert **Fig. 1c**). In addition to enrichment at TSS, footprint DNA also exhibits a prominent signal at DNase-hypersensitive sites (DHS) from a reference annotation of the B-cell line GM12878, whereas the regular cfDNA signal oscillates at neighboring genomic locations and is depleted at the DHS (**Fig. 1d**). Taken together, footprint DNA accumulates at open chromatin or TSS of genes, whereas regular cfDNA, i.e. nucleosomes, was clearly depleted in these regions.

**Fig. 1.**
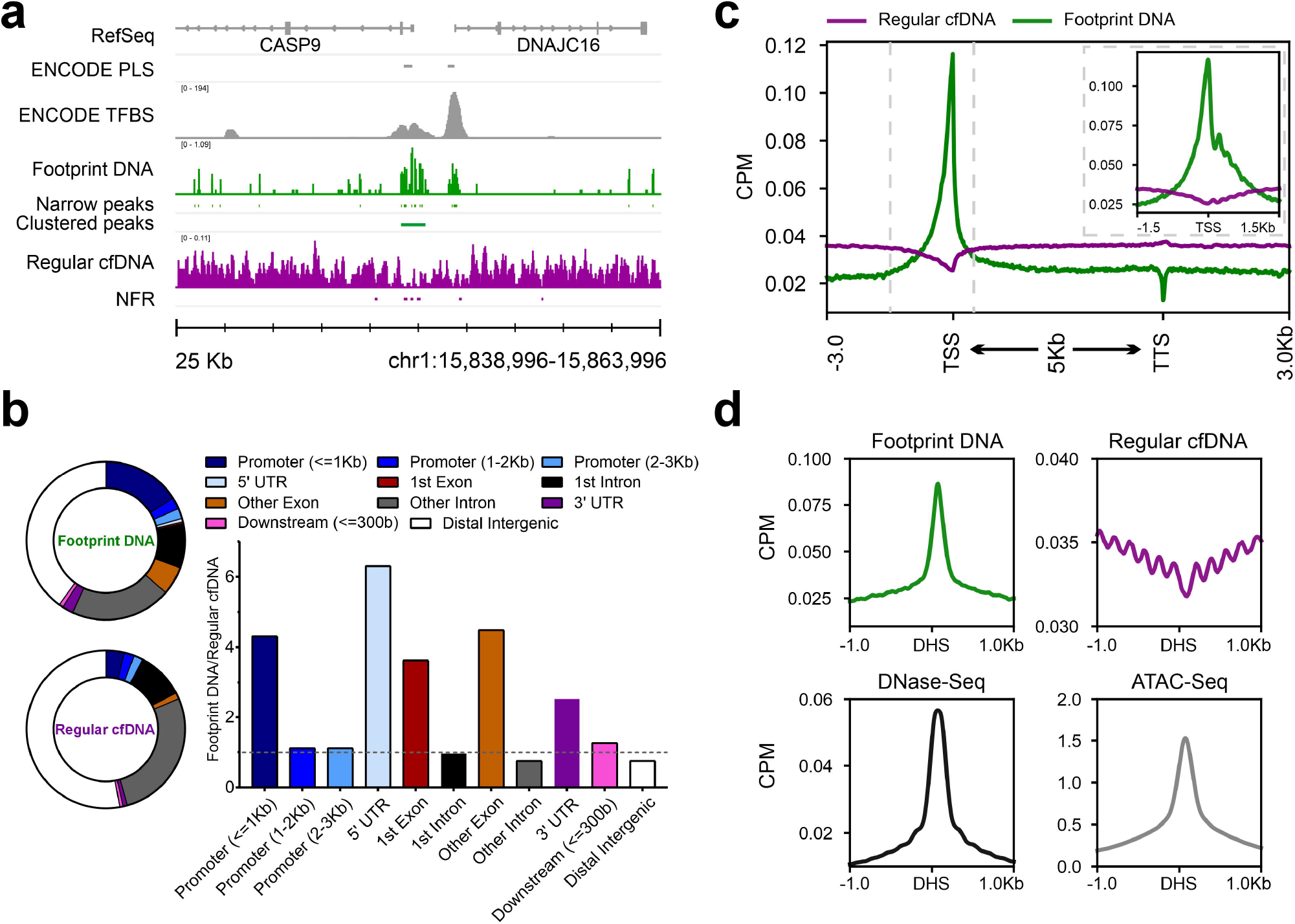
Footprint DNA is enriched in regulatory regions of genes and open chromatin. (a) Coverage profile of footprint DNA (green, S03) and regular cfDNA (violet, average of S05-S08). The footprint DNA profile shows narrow and clustered peaks, while regular cfDNA shows nucleosome-free regions (NFRs, depletion of reads). RefSeq genes, ENCODE promoter-like structures (PLS), and ENCODE transcription factor binding sites (TFBS) based on ChIP-Seq experiments are included as references. (b) Pie charts display the proportions of annotated genomic features for clustered peaks from footprint DNA (S03) and NFRs from regular cfDNA (S05-S08 merged). The bar plot shows the ratio between clustered peaks and NFRs for each genomic feature. (c) Average coverage profiles for footprint DNA (S03) and regular cfDNA (S06) along all annotated protein-coding genes. The gene bodies of all genes are scaled to 5 kilobases. Dashed vertical lines indicate the interval shown in the subplot. The subplot shows average profiles at the transcription start site without re-scaling of the gene body. (d) Average coverage profiles for footprint DNA (S03), regular cfDNA (S06), and publicly available ATAC-seq data at DNase hypersensitive sites (DHS) derived from publicly available DNase-seq data.

### Liquid footprinting detects binding of transcription factors

Since footprint DNA could be protected from nuclease digestion by binding to regulatory DBPs, peak regions were examined for encompassing transcription factor binding motifs. Accordingly, a consensus peak set was defined from peaks of liquid footprint sequencing data of four healthy individuals. Transcription factor motif enrichment analysis at the genomic locations of these consensus peaks revealed a significant enrichment of 203 transcription factor binding motifs out of 401 listed in the reference motif database HOCOMOCO. The 203 transcription factor motifs belong to 46 transcription factor families from nine transcription factor superclasses. ChIP-Seq validated transcription factor binding sites from ENCODE3 reveal clear enrichment signals for Nuclear Factor Erythroid 2 (NFE2), RE1 Silencing Transcription Factor (REST), and Spi-1 Proto-Oncogene (SPI1) in footprint DNA, for example. On the other hand, regular cfDNA is depleted at these binding sites. Overall, the average profile of all ChIP-seq based transcription factor binding sites (TFBS) shows a clear enrichment of footprint DNA in contrast to regular cfDNA independent of the transcription factor (**Fig. 2a**). ChIP-Seq validated binding sites of CCCTC-Binding Factor (CTCF) show an even more pronounced signal for footprint DNA, while regular cfDNA exhibits a high-frequency oscillation occupancy adjacent to these binding sites (**Fig. 2b**). In good agreement, DNase-seq data shows a combination of a footprint peak with adjacent oscillation patterns, while ATAC-Seq detects a peak representative for open chromatin at the CTCF binding sites (**Fig. 2b**). Our data suggest that footprint DNA most specifically reveals protection of DNA through DBPs at regulatory sites with high resolution as exemplified by CTCF, NFE2, REST, or SPI1 (**Fig. 2c**). Previously it has been shown that a significant fraction of short cfDNA exists as short single-stranded DNA (ssDNA). In order to analyze how short ssDNA signals compare to footprint DNA signals, we examined transcription start sites and transcription factor binding sites with data from Snyder *et al*. for short cfDNA sequencing (single-stranded library preparation)^6^. We found a superior signal-to-noise ratio for DBP footprints, allowing the detection of signals, e.g. TP53BP1, that were not detectable in short ssDNA of Snyder *et al*. (**Supplementary Fig. S6**)^6^. These data suggest that footprint DNA is not a degradation product of regular cfDNA but rather represents a biological entity of its own that captures DBP footprints.

**Fig. 2.**
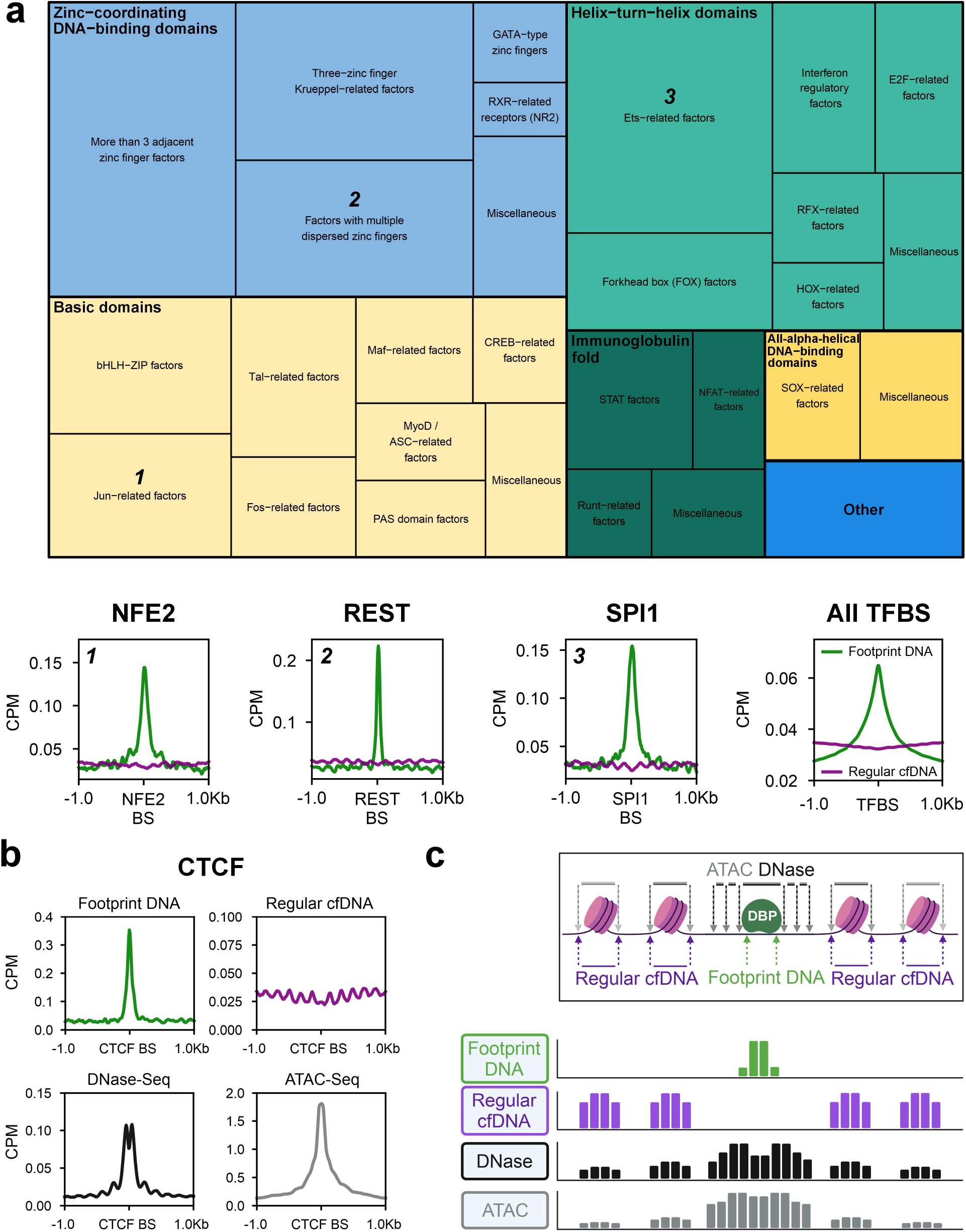
Liquid Footprinting captures the protection of the genome through various transcription factors in a direct mechanism. (a) Treemap showing transcription factors whose DNA motif was significantly enriched in footprint DNA consensus peaks from four healthy individuals. Over 200 transcription factor motifs of all nine transcription factor superclasses in the database were identified as enriched in footprint DNA peaks. The size of squares in the treemap encodes the relative number of transcription factor motifs per class (thin outlined boxes) and superclass (thick outlined boxes). As examples, average coverage profiles of footprint DNA (S03) and regular cfDNA (S06) are shown for three transcription factors (NFE2, REST, and SPI1) from three different transcription factor superclasses. Average profiles are based on the 1000 most robust binding sites annotated in the Gene Transcription Regulation Database (GTRD). In addition, the average profile of all ChIP-seq TFBS annotated in ENCODE3 is shown for footprint DNA in comparison to regular cfDNA.(b)Average coverage profile of footprint DNA (S03), regular cfDNA (S06), DNase-seq, and ATAC-seq at CTCF binding sites (CTCF BS). (c) Inferred molecular model for the formation of footprint DNA at the binding site of a DNA-binding protein (DBP) surrounded by two nucleosomes on either side. Arrows indicate the endpoints of DNA fragments for the respective sequencing technique. The resulting theoretical coverage tracks are depicted for each sequencing technique.

### Correlation to epigenetic activation and gene transcription

Given that footprint DNA fragments are likely derived from the protection of regulatory DBPs and are overrepresented at open chromatin regions we investigated a potential relationship between footprint DNA signal strength (i.e. local read enrichment) and promoter activation state. Annotated promoter-like structures were classified as active or inactive promoters based on cell-free histone 3 lysine 4 triple-methylation (H3K4me3) signals from publicly available ChIP-Seq data of a healthy individual^9^. Active promoters with strong H3K4me3 signal showed a markedly higher coverage of footprint DNA than promoters with weak or no H3K4me3 signal, whereas regular cfDNA shows exactly the opposite behavior (**Fig. 3a**). Moreover, the strongest signals for H3K4me3, representing the nucleosome positions, occur at local minima of footprint DNA. DNA methylation is known to be an essential regulator of gene activity and is associated with transcription factor binding and thus potentially DNA footprint signals. Consequently, we classified CpG islands (CpGi) into methylated and unmethylated CpGi based on the signal strength of cell-free Methyl-CpG-Binding Domain Sequencing (cfMBD-Seq) data from a healthy individual. Strongly methylated CpGi show a weaker accumulation of footprint DNA compared to unmethylated CpGi, while regular cfDNA again behaves the opposite (**Fig. 3b**), demonstrating the relationship between DNA methylation and footprint DNA. To further analyze a connection between localized footprint DNA signals and downstream gene transcription we used publicly available cell-free RNA-seq data from five healthy individuals^10^. Based on the average transcript abundance level of protein-coding genes, four subsets of genes were defined: ‘no expression’, ‘low expression’, ‘medium expression’, and ‘high expression’ genes (**Fig. 3c**). Genes with medium or high expression show considerable enrichment of footprint DNA reads at their respective TSS, whereas genes with low or no expression show no substantial enrichment. Regular cfDNA again behaves contrary to footprint DNA and shows a much less pronounced difference between active expression (high and medium) and low - or no expression (**Fig. 3d**). Consistent with the definition of these gene subgroups, H3K4me3 histone modifications increase and DNA methylation levels decrease as expression levels increase at the respective TSS of the genes (**Supplementary Fig. S7b and c**). For a more detailed analysis of the joint effect of DNA footprint signals and DNA methylation in regulatory elements of genes on its transcription, we used matched sequencing data from four septic patients. Different combinations of liquid footprints and DNA methylation signals have profound effects on the gene expression level distribution (**Fig. 3e**). Genes with low DNA methylation in their proximal CpG island and a high liquid footprint at their core promoter are likely to have high transcription levels (clusters 2, 4, and 6), while genes with high DNA methylation and a low liquid footprint are more likely to have low transcription levels (clusters 8 and 9). A transition between the activating - and repressing regulatory modes with mixed liquid footprint and DNA methylation signals gives rise to mixed gene transcription levels (clusters 3, 5, 7, and 10; **Fig. 3f**). In summary, the signal strength of footprint DNA is higher in active promoters than in inactive promoters, higher in unmethylated CpGi than in methylated CpGi, and higher in TSS of actively transcribed genes than in TSS of untranscribed genes. Thus, footprint DNA is enriched at loci with markers for epigenetic activation and transcriptionally active genomic regions.

**Fig. 3.**
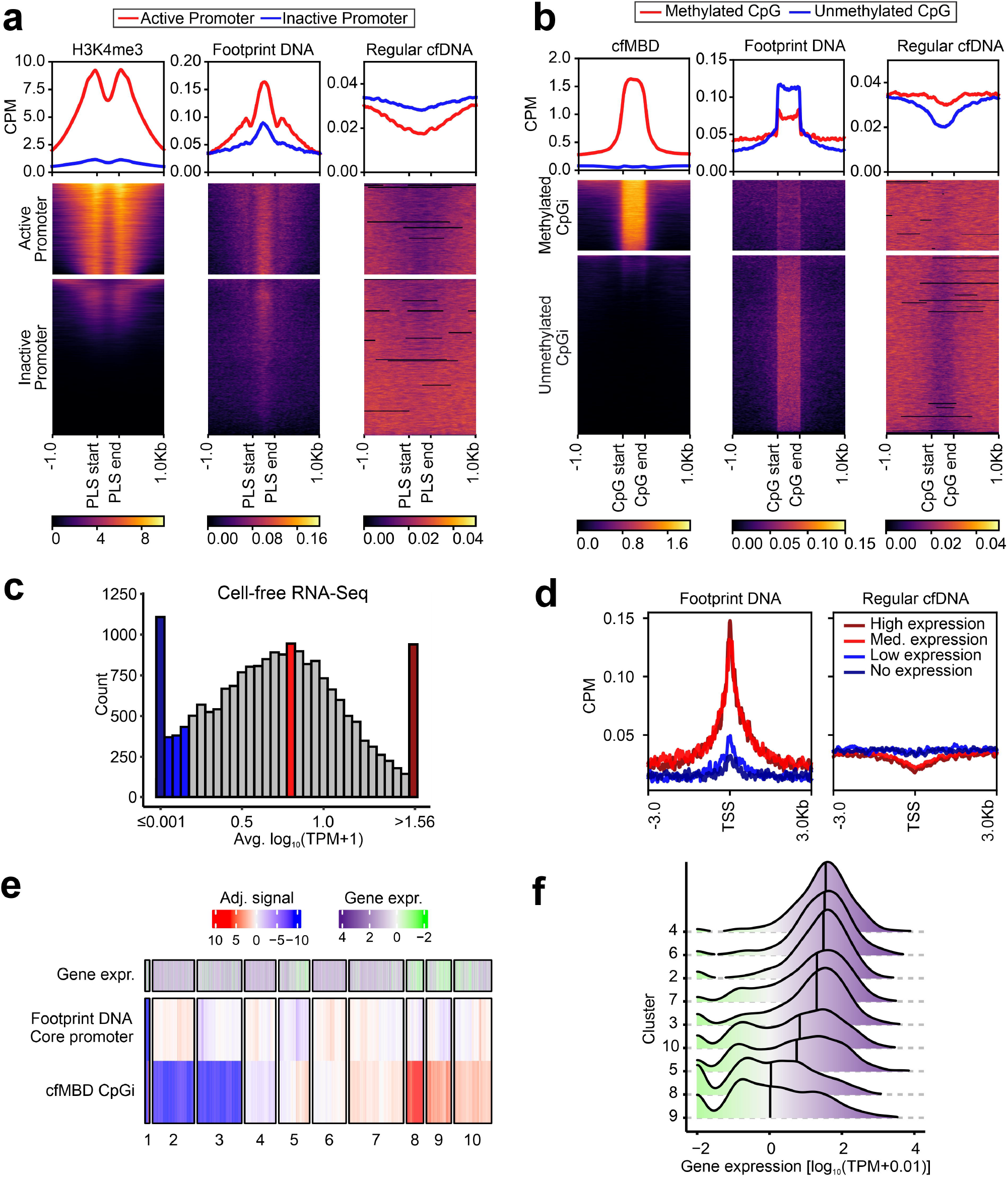
Footprint DNA is enriched at loci with markers of epigentic activation and transcriptionally active regions. (a) Annotated promoter-like structures (PLS) from ENCODE are clustered based on publicly available cell-free H3K4me3 ChIP-Seq signal strength into two clusters, representing active (red) - or inactive (blue) promoters. Average coverage profiles for footprint DNA and regular cfDNA at active or inactive promoters demonstrate the influence of the promoter activation status. (b) Annotated CpG islands are clustered based on cell-free Methyl-CpG-Binding Domain (cfMBD) sequencing signal strength into two clusters. Average coverage profiles for footprint DNA and regular cfDNA at methylated - or unmethylated CpG islands reveal the influence of methylation levels at CpG islands. (c) Histogram showing average expression levels of protein-coding genes in publicly available cell-free RNA sequencing data. For each category 5 % of all analyzed genes were selected (938 genes each for: no expression (dark blue), low expression (blue), medium expression (red), and high expression (dark red). (d) Average coverage profiles for footprint DNA and regular cfDNA at the transcription start sites of the defined gene groups of (c) reveal the influence of transcriptional activity. All data in (a) – (d) were generated from samples of healthy individuals. S03 was used as the footprint DNA dataset and S06 was used as the regular cfDNA dataset. (e) Clustered heatmap with average footprint DNA signals at core promoters and average cfMBD signals at CpG islands of protein coding genes. Average gene expression levels from whole blood are annotated. (f) Sorted ridgeplot of the gene expression annotation per cluster from (e) shows the composite influence of liquid footprints and DNA methylation on gene expression level distributions. Data in (e) and (f) were created from samples S26-S37.

### Disease-specific footprint DNA signatures in liquid biopsies

In order to identify disease-specific signatures, footprint DNA data were generated for four biological replicates of four different clinical indications: two types of gastrointestinal carcinomas (PDAC and CRC), as well as sepsis and Post-OP controls (**Supplementary Table S1**). Post-OP patients were selected as controls because they show cfDNA concentration levels comparable to septic patients. Comparison of consensus peak sets for PDAC vs. CRC revealed 731 differentially enriched loci (**Fig. 4a and Supplementary Fig. S8**) and 1,107 loci with a differential enrichment when comparing sepsis with Post-OP (**Fig. 4b and Supplementary Fig. S8**). Principal component analysis (PCA) based on all differentially enriched regions (DERs) demonstrated a clear separation of all four clinical indications by the first two principal components (**Fig. 4c**). Two exemplary DERs demonstrate a distinct differential DNA footprint near a TSS in the context of a larger locus with transcription factor binding sites (**Fig. 4d and e**). A differentially enriched transcription factor binding site was detected in the promoter of the lymphocyte cytoprotein 1 (*LCP1*) gene in CRC patients compared to PDAC patients (**Fig. 4d**), whereas another protected transcription factor binding site was detected near the promoter of the Plexin C1 gene (*PLXNC1*) in sepsis patients in contrast to Post-OP patients (**Fig. 4e**). In addition to differential analysis of signal strength at defined consensus peaks, differential enrichment of transcription factor binding motifs was also analyzed in all consensus peaks. For this purpose, the relative abundance of transcription factor motifs in the consensus peaks of one condition was compared with consensus peaks of the respective condition to be compared. Enrichment of 14 different transcription factor motifs was detected for PDAC in comparison to CRC, 19 for CRC in comparison to PDAC, 14 for sepsis in comparison to Post-OP, and 126 for Post-OP in comparison to sepsis (E-value ≤ 10, **Fig. 4f-i**). The binding motif with the strongest enrichment in PDAC patients compared to CRC patients belongs to the Recombination Signal Binding Protein For Immunoglobulin Kappa J Region (RBPJ; **Fig. 4f**), for example. In this context, one of the physiological roles of the transcription factor RBPJ is the regulation of early pancreatic cell development. Overall, liquid footprinting enables the detection of condition-specific enrichment signatures in liquid biopsies, which can be used to discriminate different diseases for diagnostic purposes. In addition, footprint DNA sequencing might also help identify transcription factors that may have physiological relevance to the condition.

**Fig. 4.**
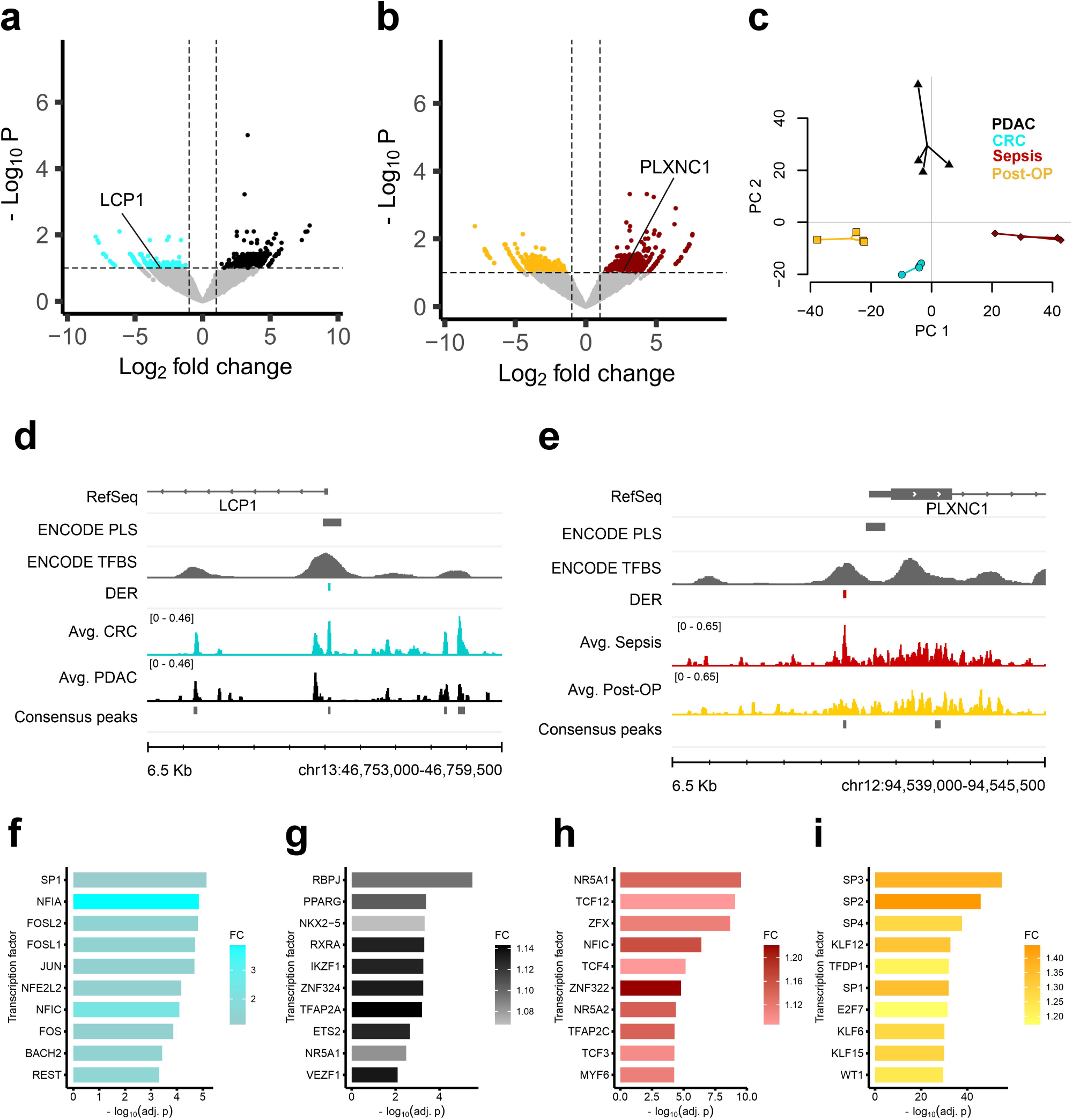
Comparison of liquid footprint data reveals condition-specific protection of transcription factor binding sites and transcription factors. (a) Differential enrichment analysis for pancreatic ductal adenocarcinoma (PDAC) samples with colorectal cancer (CRC) samples identifies differentially enriched transcription factor binding sites (Adj. p-value ≤ 0.1 and |FC| ≥ 2). (b) Differential enrichment analysis for sepsis samples with post-operative (Post-OP) samples identifies differentially enriched transcription factor binding sites (Adj. p-value ≤ 0.1 and |FC| ≥ 2). (c) Principal component analysis based on identified differential transcription factor binding sites separates all conditions. Four biological replicates were used per condition. In addition to the samples, a centroid for each group is depicted as a larger data point. Variance explained: PC1 = 29.6 %, PC2 = 20.9 %. (d) Example for a differentially enriched TFBS in CRC with novel TF binding in the promoter of LCP1. (e) Example for a differentially enriched TFBS in sepsis with novel TF binding in the promoter of PLXNC1. (f) - (i) Top 10 differentially enriched TF motifs in: (f) CRC consensus peaks in comparison to PDAC consensus peaks. CTCF and CTCFL were identified as well but were not included in the figure. (g) PDAC consensus peaks in comparison to CRC consensus peaks. (h) Sepsis consensus peaks in comparison to Post-OP consensus peaks. (i) Post-OP consensus peaks in comparison to sepsis consensus peaks. CTCF and CTCFL were identified as well but were not included in the figure.

## Discussion

In this study, we present a novel approach for comprehensive DNA footprinting in liquid biopsies by analysis of short double-stranded plasma cfDNA. Our liquid footprinting approach comprises preparative gel electrophoresis to specifically enrich cfDNA fragments with a mean length of ∼40 bp in combination with high throughput next-generation sequencing. We observed a strong enrichment of short footprint DNA at the TSS of genes, where regular cfDNA is depleted. A closer inspection of the average signals at the TSS also showed an oscillation pattern reciprocal to that of regular cfDNA at the 5’UTR. These findings indicate a regular shift of nucleosomes from the promoter to the 5’UTR of genes and protection by regulatory DBPs between displaced nucleosomes as revealed by footprint DNA enrichment. In line with the enrichment of footprint DNA at TSS we also found that footprint DNA fragments showed a higher GC content than the human genome on average (footprint DNA = 57.8 %, human genome = 40.9 %). Gene regulatory elements in humans possess an increased GC content. Consequently, footprint DNA fragments and transcription factor binding sites that are enriched in such regions of the genome, have elevated GC contents. Footprint DNA also accumulates at DNase hypersensitive sites. At these open chromatin locations, regular cfDNA exhibits a combined signal originating from high-frequency and low-frequency nucleosomal oscillation signals, as described by Ulz *et al*.^11^. Comparable to this finding, we observed two different types of footprint DNA signals created by DBPs with a narrow binding signal, such as CTCF, and multiple adjacent protein binding events that result in a broader signal. The observed narrow signal of regulatory proteins, like CTCF, SPI1, or REST can be explained by their ability to initiate nucleosome displacement in closed chromatin (pioneer factors)^12–15^. Pioneer factors can bind directly to closed chromatin without the need or presence of auxiliary proteins. In contrast, the binding sites of regulatory proteins such as MYC or TP53BP1 exhibit a much broader signal. These regulatory proteins bind in open chromatin, accompanied by the presence of additional regulatory proteins. Consequently, we also found a striking correlation between DNA footprint signals and markers representative for transcriptional regulation. In this context, we determined that promoters with H3K4me3 histone modifications, a characteristic for active gene transcription, revealed stronger footprint DNA signals than those without. Furthermore, we observed a opposing correlation between methylation levels of CpGis and corresponding enrichment of footprint DNA. Since unmethylated CpGi are considered markers of epigenetic activation, depletion of regular cfDNA and the presence of DNA footprinting signals indicate a protection from regulatory activating factors at such loci. Regarding gene expression, footprint DNA signals at the TSS are increased at actively transcribed genes compared to genes with low expression. However, footprint DNA does not strictly capture continuous dynamics of gene expression levels, rather more it exhibits a binary switch between high and low expression. Overall, footprint DNA is correlated with DNA-methylation signatures, H3K4me3 histone modifications, and transcriptional activity of downstream gene loci. Remarkably, we found consistent enrichment of footprint DNA where regular cfDNA is depleted. We propose that the molecular origin of footprint DNA most likely is double-stranded cfDNA which has been protected from enzymatic digestion by regulatory proteins, including transcription factors. In agreement with this model, we found that footprint DNA consensus peaks comprise a plethora of transcription factor motifs and that known transcription factor binding sites revealed clear enrichment for footprint DNA. Taken together, our data provide strong evidence that footprint DNA represents a characteristic subset of cfDNA that is distinct from nucleosomal cfDNA and therefore most likely does not originate from the degradation of regular cfDNA.

In contrast to footprint DNA, which is characterized by short double-stranded DNA (short dsDNA), publications by Snyder *et al*., Hudecova *et al*., and Hisano *et al*. reported on the presence of short ssDNA^6,16,17^. Contrary to short dsDNA, short ssDNA fragments may represent a significant fraction of total cfDNA comprising as much as 20% of a sequencing library. However, short ssDNA demonstrated that the relative signal strength at transcription start sites or TFBS of proteins, e.g. MYC, is very low compared to short dsDNA. Therefore, we assume that cfDNA originally protected by regulatory DBPs is short dsDNA whereas short ssDNA might be of other biological origin with higher signal-to-noise ratios. Hudecova *et al*., and Hisano *et al*. suggested that much of the short ssDNA could originate from non-canonical DNA structures, such as G4 quadruplexes, which may largely superimpose signals from regulatory protein DNA-binding events.

Transcription factor motifs from footprint DNA revealed sets of differentially enriched transcription factors between different conditions including samples from septic, colorectal carcinoma (CRC) as well as pancreatic cancer (PDAC) patients, suggesting a diagnostic potential for liquid biopsy applications. Sepsis is characterized by a complex interplay of pro-inflammatory and anti-inflammatory processes orchestrated by regulators of the immune system. Seven out of the top 10 differentially enriched transcription factor motifs identified in septic samples are linked to regulatory pathways of the immune system. For example, transcription factor 4 (TCF4, also known as E2-2 or ITF-2) regulates genes for the differentiation of dendritic cells into interferon-producing plasmacytoid dendritic cells^18^. TCF4 also regulates the immune response as a downstream target of TLR2 signaling to induce the expression of immunoregulatory genes such as interleukin 10 (IL-10)^19^. The identified transcription factor ZFX is known to be involved in the maintenance of peripheral T cells as well as expansion and maintenance during B cell development and peripheral homeostasis^20,21^. For CRC patients, specificity protein 1 (SP1) was identified as the top hit, and several Fos/Jun family transcription factors were identified as differentially enriched. SP1 is a ubiquitous transcription factor and mediator of critical physiological pathways including cell cycle, proliferation, and metastasis. SP1 plays a key role in regulating genes involved in CRC growth and metastasis^22^. Two members of the Fos/Jun family, FOS Like 1 (FOSL1) and FOS Like 2 (FOSL2) are known to promote tumorigenesis and metastasis in colon cancer^23,24^. For PDAC patients, Recombination Signal Binding Protein For Immunoglobulin Kappa J Region (RBPJ) was identified as the corresponding top hit. RBPJ is known to form a heterocomplex with Pancreas Associated Transcription Factor 1a (PTF1A) and their interaction is required in the early stage of pancreatic growth, morphogenesis, and lineage fate decision^25^. In addition, Peroxisome proliferator-activated receptor gamma (PPARγ) and Retinoid X receptor alpha (RXRα) were identified, which are known to form a heterocomplex^26^. PPAR γ is a key regulator of adipocyte differentiation, regulates insulin and adipokine production and secretion, and is associated with PDAC. In addition, RXRα promotes proliferation and inhibits apoptosis of pancreatic cancer cells^27^. Moreover, we found disease-specific differentially enriched loci that may enable clear separation of patients with PCA. For example, in CRC patients, we found differential protection from DBPs near the TSS of the lymphocyte cytosolic protein 1 (*LCP1* also known as *plastin-2*) gene compared with PDAC patients. *LCP1* is associated with CRC progression, prognosis, and metastasis, and patients with late-stage CRC have higher expression levels of plastin-2^28–30^. In septic patients, we found a strong signal of footprint DNA near the TSS of the Plexin C1 gene (*PLXNC1*) compared with post-operative controls. PLXNC1 augments adhesion, transmigration, and activation of neutrophils during acute lung injury and is induced during an acute inflammatory response^31,32^.

Taken together, analysis of short double-stranded cfDNA might provide the most accurate picture of a genome-wide transcription factor footprint in liquid biopsies. With the ability to identify disease-specific transcription factor binding site enrichment for patient classification, we see considerable potential in the application of footprint DNA sequencing for liquid biopsy applications.

The results presented in this study are based on comparably small sample sizes. Future validation studies with larger cohort sizes will be useful for validation. In addition to validity and utility, cost-effectiveness is an essential factor for clinical diagnostic methods. Although our workflow already represents a resource-efficient alternative to ultra-deep high-throughput sequencing of total cfDNA, a targeted and even more economical approach might be advantageous for implementation. With the new liquid footprinting approach, that directly maps the protection across the genome via regulatory proteins, we want to add a new tool to liquid biopsy diagnostics that could improve the detection of cancer or enable clinically relevant differential diagnosis of cancer types.

## Methods

### Blood samples

This study included blood samples from nine healthy individuals, four individuals with pancreatic ductal adenocarcinoma, four individuals with colorectal carcinoma, four individuals with sepsis, and four individuals that underwent major abdominal surgery (**Supplementary Table S1**). Blood from healthy individuals was acquired commercially from Biomex (Heidelberg, Germany). Septic patients participated in a previously published, prospective observational clinical study which was conducted in the surgical intensive care unit of Heidelberg University Hospital, Germany between November 2013 and January 2015 (S13-S16 and S25-S37; German Clinical Trials Register: DRKS00005463)^33^. Patients without cancer (non-septic controls, i.e. Post-OP) that underwent major abdominal surgery and all individuals with cancer were recruited at the University Hospital of Erlangen with the approval of the local ethics committee and the clinical trial number 180_19 B (S09-S12 and S17-S24). All experiments were performed in accordance with the study protocol approved by the Ethics Committee.

### Cell-free DNA isolation

Plasma was prepared by centrifugation of whole blood for 10 min at 1,600 g and 4 °C. Afterward, blood plasma was centrifuged again for 10 min at 16,000 g and 4 °C. Afterward, 1.1 mL of the supernatant was transferred into a fresh 1.5 mL DNA LoBind tube and stored at -80 °C. If necessary, the sample volume was filled up with freshly prepared 1 x phosphate-buffered saline solution. Cell-free DNA was isolated with the QIAsymphony SP DNA Preparation System and the QIAsymphony DSP Circulating DNA Kit (Qiagen, Hilden, Germany) according to the manufacturer’s advice. Eluted cfDNA was quantified with the Qubit dsDNA HS Assay Kit (Thermo Fisher, Waltham, Massachusetts, USA) and the cfDNA quality was assessed by the Fragment Analyzer High Sensitivity DNA Kit (Agilent, Santa Clara, California, USA).

### Next-Generation sequencing

Sequencing libraries for regular cfDNA were prepared with the NEBNext Ultra II DNA Library Prep Kit (New England Biolabs, Ipswich, Massachusetts, USA) according to the manufacturer’s protocol. 0.5 ng isolated cfDNA was used as input. The NEBNext Adaptor was diluted 1:25 for all reactions. PCR was performed with 10 PCR cycles and 4 μL primers. Sequencing was performed on a NextSeq 2000 (Illumina, San Diego, California, USA) with 100 bp single end reagent kits.

Sequencing libraries for short double-stranded cfDNA (footprint DNA) were prepared from 3 ng to 15 ng of cfDNA, depending on the clinical condition, using the NEXTFLEX Cell free DNA-Seq Kit (V2) (Perkin Elmer, Pittsburgh, Pennsylvania, USA) according to the manufacturer’s advice with one exception: the final library after PCR amplification was eluted in 20 μL nuclease-free water. Library generation was performed with the Biomek FXP workstation (Beckman Coulter, Brea, California, USA). Library quality was assessed by the Fragment Analyzer High Sensitivity DNA Kit (Agilent, Santa Clara, California, USA) and the concentration was measured by the Qubit dsDNA HS Assay Kit (Thermo Fisher, Waltham, Massachusetts, USA). Size selection of cfDNA libraries was performed using a BluePippin instrument (Sage Science, Beverly, Massachusetts, USA). To select the footprint DNA portion in the range of 150-200 bp, samples were applied to a 3% agarose BluePippin cassette according to the manufacturer’s protocol. Briefly described, samples were filled up with water to 30 μL and were mixed with 10 μL of supplied internal marker (100 bp to 250 bp). 23 μL of the eluted, size-selected sample was reamplified with 25 μL of NEXTFLEX PCR Master Mix 2.0 and 2 μL of 1:10 diluted NEXTFLEX Primer Mix 2.0 as described in the NEXTFLEX Cell free DNA-Seq Kit (V2) (Perkin Elmer, Pittsburgh, Pennsylvania, USA), step C PCR amplification. Afterward, samples were purified with 1.2 x the volume of AMPureXP beads (Beckman Coulter, Brea, California, USA) according to the manufacturer’s advice. Size selection performance was evaluated by the Fragment Analyzer High Sensitivity DNA Kit (Agilent, Santa Clara, California, USA). If the sample still contained fragments outside the target range of 150 bp to 200 bp, size selection was repeated as previously described, with the exception that the input sample was adjusted to 30 μL with water and the reamplification step was performed with 5 to 8 cycles. After size selection, sequencing was performed on a NextSeq 2000 (Illumina, San Diego, California, USA) with 100 bp single end reagent kits.

Methylation enrichment of cfDNA was performed using the EpiMark Methylated DNA Enrichment Kit (New England Biolabs, Ipswich, Massachusetts, USA) according to the manufacturer’s instructions. Two reactions with 5-15 ng of cfDNA input each were performed in parallel per sample. Methylated cfDNA was eluted sequentially from both reactions using the same 50 μL of nuclease-free water to increase yield. Sequencing libraries were prepared with the NEBNext Ultra II DNA Library Prep Kit (New England Biolabs, Ipswich, Massachusetts, USA) according to the manufacturer’s protocol. 50 μL methylated cfDNA from the EpiMark procedure was used as input. The NEBNext Adaptor was diluted 1:25 for all reactions. PCR was performed with 17 PCR cycles and 3 μL primers. Sequencing was performed on a NextSeq 2000 (Illumina, San Diego, California, USA) with 100 bp single end reagent kits.

For whole blood RNA-Seq, blood of patients was collected in PAXgene Blood RNA tubes (BD, Heidelberg, Germany), incubated at room temperature for 2 h to achieve complete lysis of blood cells, and frozen at -80 °C until further processing. Before nucleic acid isolation, tubes were thawed at room temperature for 2 h. Nucleic acid isolation was performed using the QIAcube (Qiagen, Hilden, Germany) and the PAXgene blood miRNA kit according to the manufacturer’s protocol to extract gene-encoding mRNAs. Nucleic acids were eluted in 2x40 μL Buffer BR5. The quantity and quality of the isolated RNA was determined with a Qubit Fluorometer (Thermo Fisher, Waltham, Massachusetts, USA) and a Fragment Analyzer (Agilent, Santa Clara, California, USA), respectively. Library preparation and sequencing were performed using 250 ng RNA with the ScriptSeq kit v2 (Illumina, San Diego, CA, USA). Sequencing of the libraries was performed with HiSeq2500 (Illumina, San Diego, CA, USA), with 100 bp single end reagent kits.

All acquired raw next-generation sequencing data of this study are deposited in the sequence read archive (SRA) of the national center for biotechnology information (NCBI) under the following accession number: PRJNA1033613. All data are fully available without restriction.

### Sequencing data processing

After initial quality control of liquid footprinting raw sequencing reads with FastQC (v0.11.8), the following steps were performed sequentially to remove sequencing artifacts: (1) Removal of sequencing adapters, terminal polyG sequences (min 10 sequential G’s), and quality trimming (BBTools - bbduk.sh v38.67). (2) Removal of terminal single A nucleotide added during library preparation (BBTools - bbduk.sh v38.67). (3) Size selection of sequenced fragments allowing read lengths greater than 20 bp and smaller than 60 bp (BBTools - bbduk.sh v38.67). (4) Removal of sequencing reads with low complexity, i.e. dust scores smaller than 7 (prinseq-lite v0.20.4)^34–36^. (5) Processed reads were mapped to the human reference genome assembly GRCh37 using NextGenMap (v0.5.5) with default settings^37^. (6) Mapped reads were deduplicated with samtools rmdup (v1.9), reads in blacklisted regions were removed with bedtools intersect (v2.30.0), and reads with a MAPQ value lower than 1 were removed with samtools view (v1.9)^38–40^. These reads were converted to BigWig format for visualization and other downstream analyses with deeptools (bin size = 10, normalization = counts per million (CPM), bamCoverage v3.5.1)^41^. This workflow is graphically summarized in **Supplementary Fig. S3**.

After initial quality control of regular cfDNA raw sequencing reads with FastQC (v0.11.8), the following steps were performed^36^: (1) Removal of sequencing adapters, removal of terminal polyG sequences (min 10 sequential G’s), removal of reads shorter than 50 base pairs, and quality trimming (BBTools - bbduk.sh v38.67)^34^. (2) Processed reads were mapped to the human reference genome assembly GRCh37 using NextGenMap (v0.5.5) with default settings^37^. (3) Mapped reads were deduplicated with samtools rmdup (v1.9), and reads in blacklisted regions were removed with bedtools intersect (v2.30.0)^39,40^. Mapped reads were converted to BigWig format for visualization and other downstream analyses with deeptools (bin size = 10, normalization = counts per million (CPM), bamCoverage v3.5.1)^41^.

Cell-free Methyl-CpG-Binding Domain sequencing data were processed as regular cfDNA sequencing data.

### Public sequencing data processing

Fastq files of five publicly available cell-free RNA sequencing datasets of different healthy individuals from Zhu *et al*. were obtained from the sequencing read archive (SRA)^10^. Accession numbers and unique identifiers are listed in **Supplementary Table S2**. All samples were sequenced in paired-end end mode with a read length of 150 bp and about 10 million reads per sample on average. After initial quality control of raw sequencing reads with FastQC (v0.11.8), the following steps were performed to obtain read counts for individual genes^36^: (1) Removal of sequencing adapters, quality trimming, and removal of reads shorter than 100 base pairs (BBTools - bbduk.sh v38.67)^34^. (2) Mapping of processed reads to the human reference genome assembly GRCh37 using NextGenMap (v0.5.5) with default settings, keeping only reads with a MAPQ value greater than 2^37^. (3) Gene quantification in raw read counts using featureCounts (v2.0.1) with the UCSC Genes annotation (available at: https://hgdownload.soe.ucsc.edu/goldenPath/hg19/bigZips/genes/hg19.knownGene.gtf.gz)^42^. Only protein-coding genes on the autosomal chromosomes were included in the final readcount matrix to reduce confounding by the gender of sample donors. (4) If required, raw read counts were converted to transcripts per million (TPM) using R.

Fastq files of cell-free DNA sequencing datasets of a healthy individual from Snyder *et al*. were obtained from the sequencing read archive (SRA)^6^. Accession numbers and unique identifiers are listed in **Supplementary Table S2**. After initial quality control of raw sequencing reads with FastQC (v0.11.8), the following steps were performed: (1) Removal of sequencing adapters and quality trimming (cutadapt v4.0)^43^. (2) Mapping of trimmed reads to the human reference genome assembly GRCh37 using NextGenMap (v0.5.5) with minimum map quality of 10^37^. (3) In silico size selection of the mapped reads: 35-80 nt for footprint DNA and 120-180 nt for regular cfDNA. (4) Removal of low complexity reads and de-duplication (prinseq-lite v0.20.4)^35^. (5) Removal of reads in blacklisted regions with bedtools intersect (v2.30.0)^40^. (6) For downstream analysis and visualization BigWig files were generated with deeptools (bin size = 10, normalization = counts per million (CPM), bamCoverage v3.5.1)^41^.

ATAC-Seq and DNase-Seq data from ENCODE (see **Supplementary Table S2**) were retrieved as processed bam files and only converted to other data types, like BigWig for downstream analysis. Cell-free H3K4me3 ChIP-Seq data from Sadeh *et al*. (see **Supplementary Table S2**) were retrieved as processed bed files and only converted to other data types, like bam and BigWig for downstream analysis^9^.

### Peak calling and nucleosome-free region calling

For footprint DNA sequencing data, narrow peaks and clustered peaks were called with MACS2 callpeak (narrow: --nomodel --extsize 32 --call-summits --min-length 30 -q 0.05, clustered: --broad --nomodel --extsize 32 --max-gap 100 --min-length 500 --broad-cutoff 0.05)^44^. Consensus peaks of a condition were identified using R and defined as genomic sites where a narrow peak was identified in at least three of four samples. In addition, consensus peaks less than 31 nucleotides apart were combined. For regular cfDNA, nucleosome-free regions were called on the merged dataset of four biological replicates to increase genome-wide coverage and thus reliability. Nucleosome-free regions were identified with the R packages NucDyn, utilizing nucleR, with default settings^45^. Peaks and nucleosome-free regions were annotated to genomic features in R with the ChIPseeker package and default settings^46^.

### Average coverage profiles and heatmaps

Average coverage profiles and heatmaps were created from BigWig files with deeptools (computeMatrix, plotHeatmap, and plotProfile; v3.5.1)^41^. Different genomic reference locations were used for average coverage profiles. For **Fig. 2** and **Supplementary Fig. S2**, validated binding sites of transcription factors were retrieved from Gene Transcription Regulation Database (GTRD)^47^. Here, transcription factor binding sites were ranked according to their robustness across different cell lines and tissues, i.e., the number of tissues and cell lines in which the respective binding site was found. Only the top 1000 binding sites were used for average coverage profiles. Promoter-like structures from ENCODE were converted from GRCh38 to GRCh37 with the liftOver tool from UCSC and used as reference regions in **Fig. 3a** (available at https://www.encodeproject.org/files/ENCFF379UDA/). CpGi were used as reference regions in **Fig. 3b** (available at UCSC TableBrowser, track name = cpgIslandExt). The heatmaps in **Fig. 3a and b** were clustered in two groups using the implemented k-means clustering based on the line-wise average. Missing data in heatmaps were plotted in black.

### Composite effect of DNA methylation and DNA footprint signals on gene transcription

From four sepsis patients, matched liquid footprinting, cfMBD, and whole blood RNA sequencing data were acquired (S26-S37). For combined analysis different genomic annotations were combined. The same gene annotation as used with the cfRNA sequencing data from Zhu *et al*. was utilized^10^. A core promoter annotation for GRCh37 was retrieved from the eukaryotic promoter database (EPD, version 6, available at https://epd.expasy.org/ftp/epdnew/H_sapiens/006/Hs_EPDnew_006_hg19.bb). For each gene, only the best-matching core promoters were kept (‘_1’ flag). As CpGi, the aforementioned CpGi from UCSC were used again. CpGi were assigned to genes based on their proximity to the EPD promoter of the gene in the genome, with a maximum absolute distance of 5 Kb. With these assignments, from the roughly 20,000 protein-coding genes in the annotation, about 12,000 remained with an associated core promoter and CpG island. Counts of sequencing reads of each sequencing data type and each patient were retrieved for the curated gene bodies, core promoters, and CpG island with featureCounts (v2.0.1) and converted to TPM. From the patient replicates a mean TPM value was calculated per region and sequencing type. Additionaly, a pseudocount of 0.01 was added. For better comparability of the signals, the cfMBD-Seq data in CpGi and liquid footprinting data in core promoters were additionally adjusted for GC content. The GC content of the respective regions was extracted from GRCh37 using bedtools nuc (v2.30.0). The adjusted signal was calculated as follows: A locally weighted regression function was fitted with the mean readcount value and GC content for each annotation type (R lowess function). Each mean readcount value (observed) is divided by its fitted value (expected) to obtain an oberserved over expected (OoE) ratio. For better interpretability and visualization, the values are also log_2_-scaled. Genes with comparable composite signals in the core promoter and in the CpG island were identified by k-means clustering (k=10) and visualized in a heatmap using R and ComplexHeatmap (**Fig. 3**e). The average gene expression values were added as an annotation, but had no influence on clustering. For better visualization and comparison of the distributions of gene expression values, the RNA-Seq data were also visualized in a ridge plot in R with ggplot, according to the clusters from the heatmap and sorted by descending median value (**Fig. 3f**). Clusters with less than 200 genes were excluded from this visualization as the sample was considered too small for a representative distribution.

### Transcription factor motif enrichment analysis

Enriched transcription factor motifs were identified with the AME tool from MEME suite (-- scoring avg --method fisher; v5.4.1)^48^. Input DNA sequences were retrieved from consensus peak sets and analyzed for the enrichment of motifs listed in the HOCOMOCOv11_core_HUMAN_mono database. DNA sequences of consensus peaks from footprint DNA sequencing data of healthy individuals were compared to control sequences, generated by shuffling the letters in the input while preserving the frequencies of k-mers (--shuffle). The proportions of identified transcription factor motif classes and their respective superclasses were summarized in a treemap plot using R and the treemap package (v.2.4.3). Footprint DNA sequencing data from other than healthy states were compared with each other. The DNA sequences underlying each consensus peak set were used as control sequences, e.g. consensus peak DNA sequences from CRC as input compared to consensus peak DNA sequences from PDAC as control.

### Differential enrichment analysis

Identification of differentially enriched regions (DERs) was performed with the R package DEBrowser (v1.2.0) using the implemented edgeR method with raw read counts in the combined consensus peak sets of two compared conditions, TMM normalization, a glmLRT, and dispersion = 0^49,50^. Genes were considered as differentially expressed between two conditions (four biological replicates per condition) with an adjusted p-value smaller than 0.1 and a fold change ≤ −2 or ≥ 2. Volcano plots for the differential enrichment analysis were generated with the R package EnhancedVolcano (v1.8.0). The identified DERs of both comparisons were used for a principal component analysis (PCA), and the first three principal components were visualized in R with pca3d (v0.10.2). Samples of each condition were linked to the centroid of the respective condition with a line.

## Acknowledgments

This work was financed with internal funding of the Fraunhofer society. We also thank MDs Jens and Susanne Freitag for their valuable support.

## Author contributions

J.M. and K.S. conceptualized and designed the study. C.H., M.S., L.B. and K.G. processed blood samples, isolated cfDNA, and established the size selection method for short double-stranded cfDNA. J.M. performed bioinformatics analysis. C.A. analyzed the effect of library preparation methods on short and regular cfDNA in public data. J.M., K.S., Y.V., and A.v.H. analyzed and interpreted the sequencing data. G.W., S.D, and T.B. recruited patients and collected samples. J.M. and K.S. wrote the manuscript. All authors read and approved the final manuscript.

