## Supplementary materials for "Liquid footprinting: A novel approach for DNA footprinting using short double-stranded cell-free DNA from plasma"

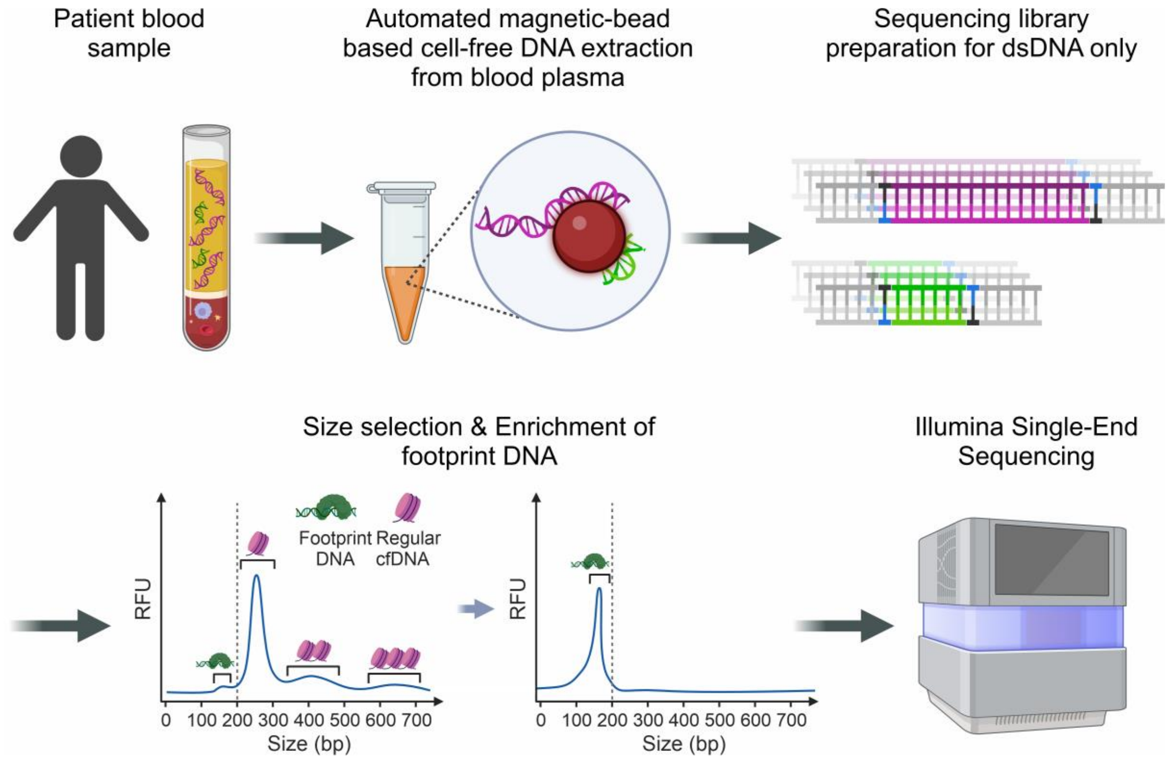


**Supplementary Figure S1 – Visual summary of footprint DNA extraction and sequencing from blood plasma.** Cell-free DNA was extracted from the blood plasma of patients using an automated magnetic bead-based kit. Sequencing library preparation was performed by fragment end-repair and adapter ligation. Only intact double-stranded DNA fragments are enriched in the final sequencing library because of the PCR amplification. Footprint DNA is enriched by size selection from sequencing libraries. Footprint DNA sequencing libraries are sequenced on an Illumina platform in single-end mode. This figure was created with biorender.

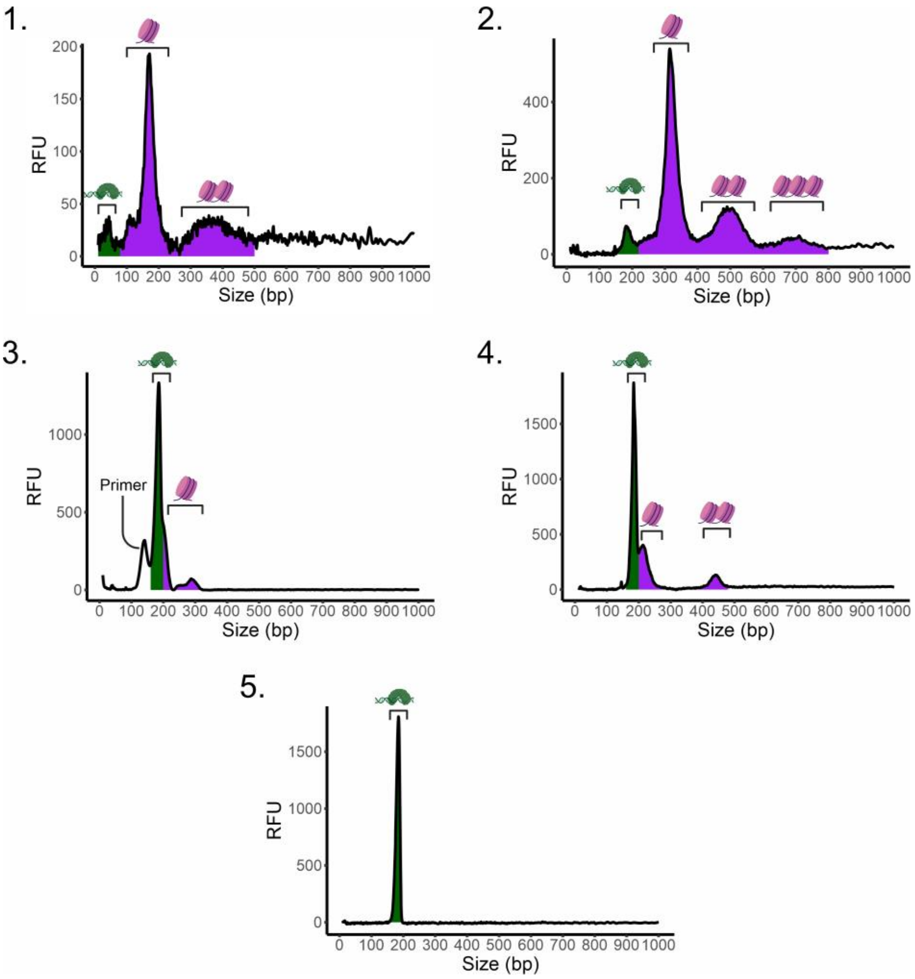


**Supplementary Figure S2 – Depiction of the full footprint DNA enrichment process from cell-free DNA for next-generation sequencing.** The steps shown correspond to: 1. Isolated cell-free DNA. 2. Double-stranded DNA sequencing library. 3. Size-selected sequencing library. 4. PCR-amplified sequencing library. 5. Sequencing library after second size selection. Size selection was performed using an automated preparative gel electrophoresis instrument. For each step, the Fragment Analyzer profiles of S19 are shown. The library preparation adds around 100 bps to the DNA fragments resulting in the fragment size shift seen from 2. onwards. The purple color indicates DNA fragments that can be assigned to nucleosomes or regular cfDNA, while the green color indicates DNA fragments that can be assigned to footprint DNA. The icons originally created with biorender for the supplementary figure S1 were reused in this figure.

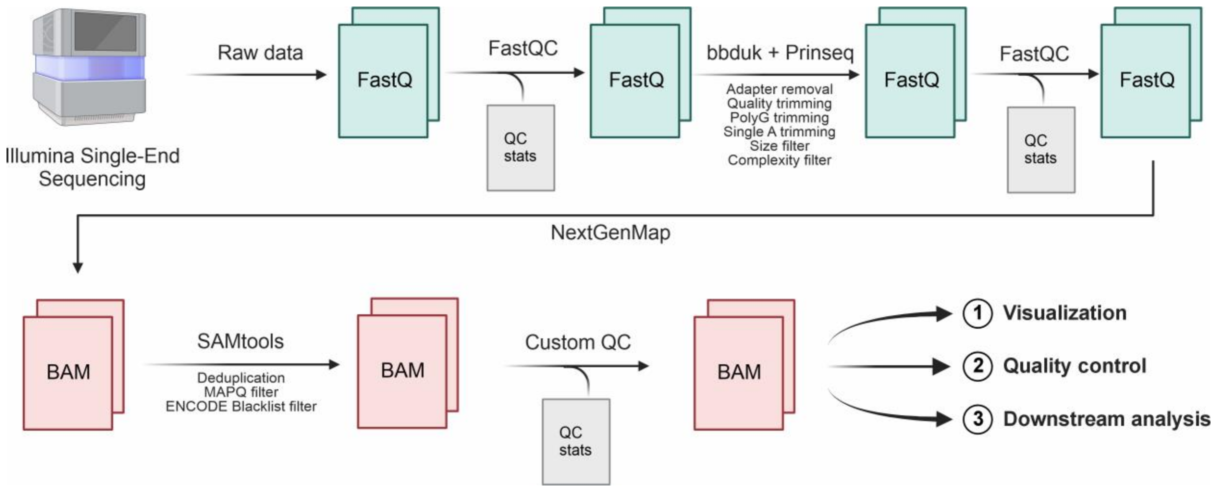


**Supplementary Figure S3 – Visual summary of the footprint DNA data processing pipeline.** In short, the raw footprint DNA sequencing data is cleaned and filtered, mapped to a human reference genome, and filtered again before further downstream analysis. The individual steps are described in detail in the methods section.

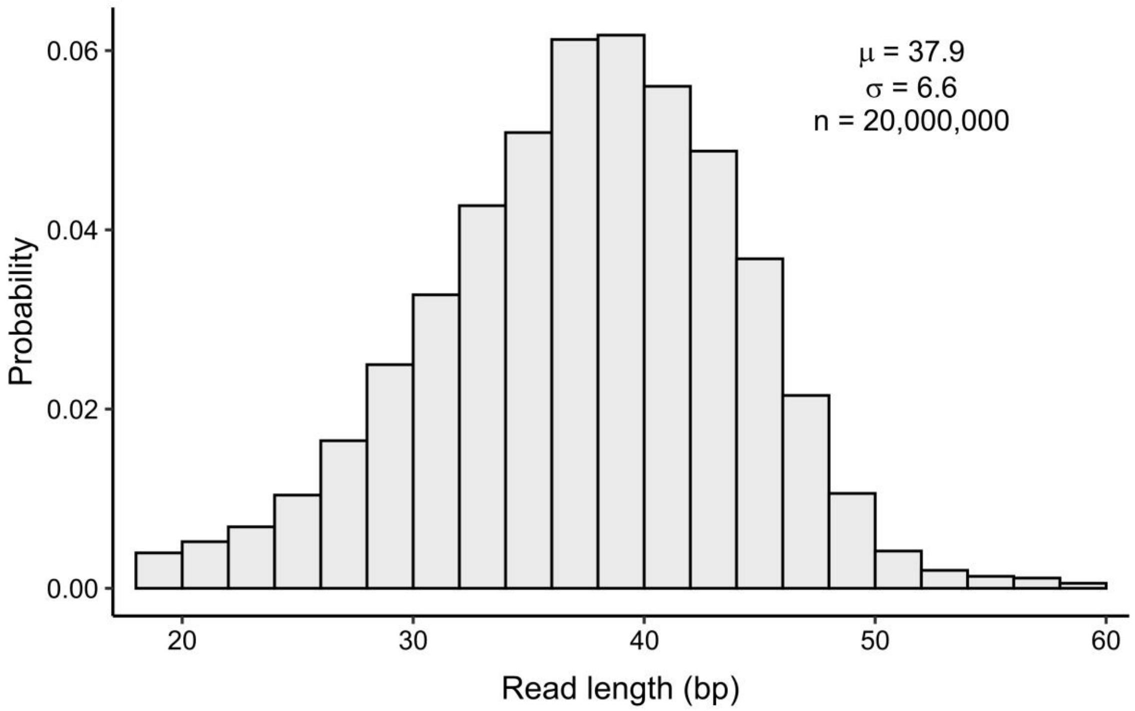


**Supplementary Figure S4 – Histogram depicting the observed length of fully processed footprint DNA sequencing reads.** One million random reads were taken from all twenty sequenced liquid footprint samples and their read lengths were plotted in a histogram (total n = 20 million). The observed distribution has a mean read length of = 37.9 bp (μ) and a standard deviation = 6.6 bps (σ).

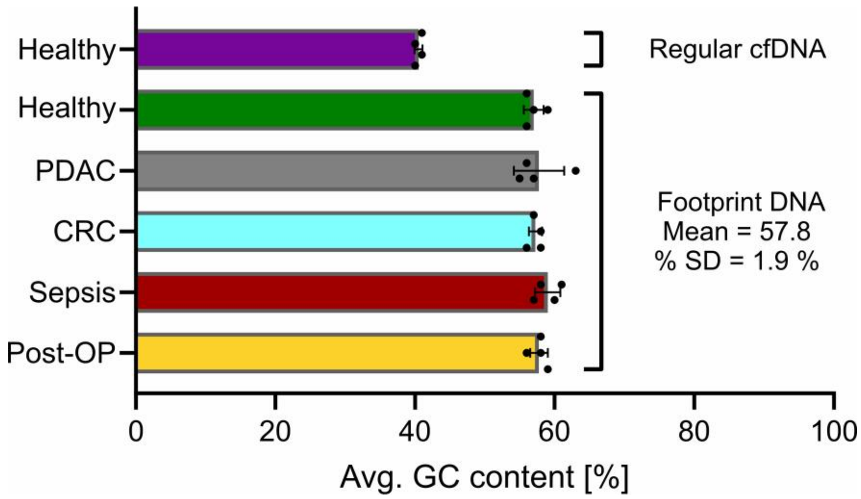


**Supplementary Figure S5 – Average GC content of processed sequencing reads for regular cfDNA and footprint DNA sequencing.** Data from the samples S01 - S24 were used for this analysis (see Supplementary Table S1). The bar lengths represent the mean value, while error bars indicate standard deviations.

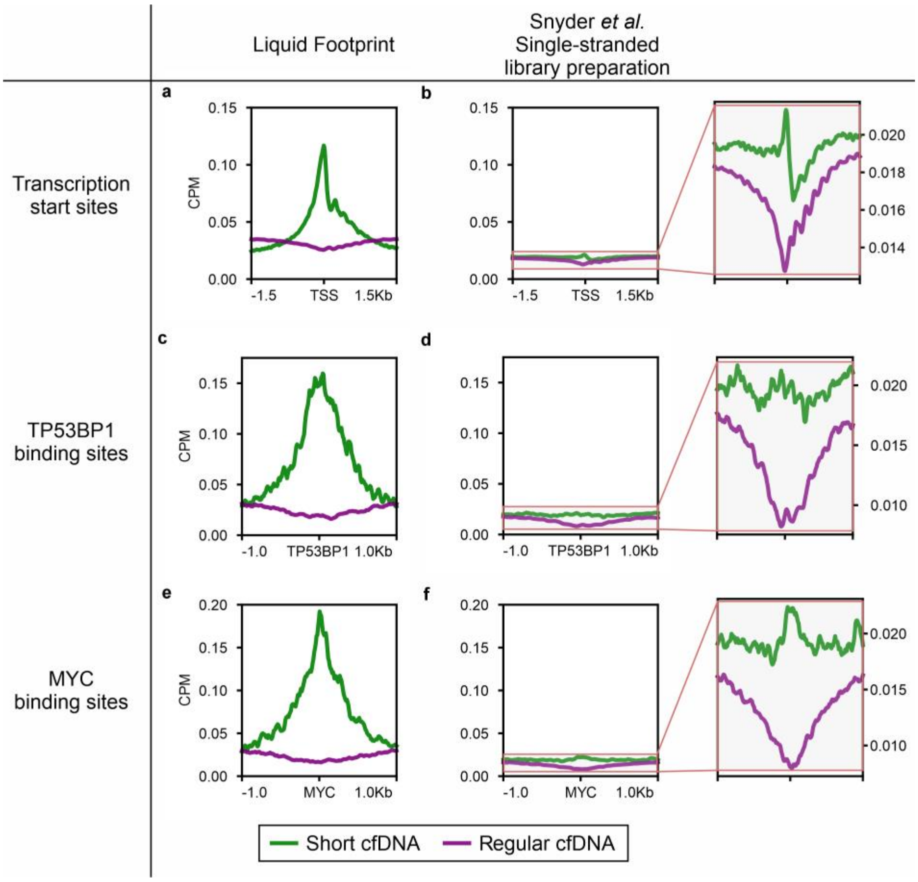


**Supplementary Figure S6 – Comparison of liquid footprinting data to single-stranded cfDNA sequencing data from Snyder *et al.*[1].** (a, c, e) Average coverage profiles based on liquid footprint sequencing data (Sequencing depths: Short cfDNA (S03) = $8.29 \cdot 10^6$, Regular cfDNA (S06) = $2.60 \cdot 10^7$). (b, d, f) Average coverage profiles based on sequencing data from Snyder *et al.* generated with a single-stranded library preparation method (Sequencing depths: Short cfDNA = $1.33 \cdot 10^8$, Regular cfDNA = $3.84 \cdot 10^8$). These average coverage profiles are also plotted on a smaller scale to better visualize the dynamics of the data. (a and b) Average coverage profiles of annotated transcription start sites for short cfDNA and regular cfDNA. (c and d) Average coverage profiles of one thousand ChIP-Seq validated TP53BP1 binding sites for short cfDNA and regular cfDNA. (e - f) Average coverage profiles of one thousand ChIP-Seq validated MYC binding sites for short cfDNA and regular cfDNA. The purple color indicates data from regular cfDNA, while the green color indicates data from short cfDNA. Ultra-deep sequencing data by Snyder *et al*. was produced from the blood plasma of a healthy individual with a single-stranded library preparation method[1]. Raw sequencing data were retrieved from SRA (file ID = SRR2130051) and split into short cfDNA (35-80 nt) and regular cfDNA (120-180 nt) *in silico*.

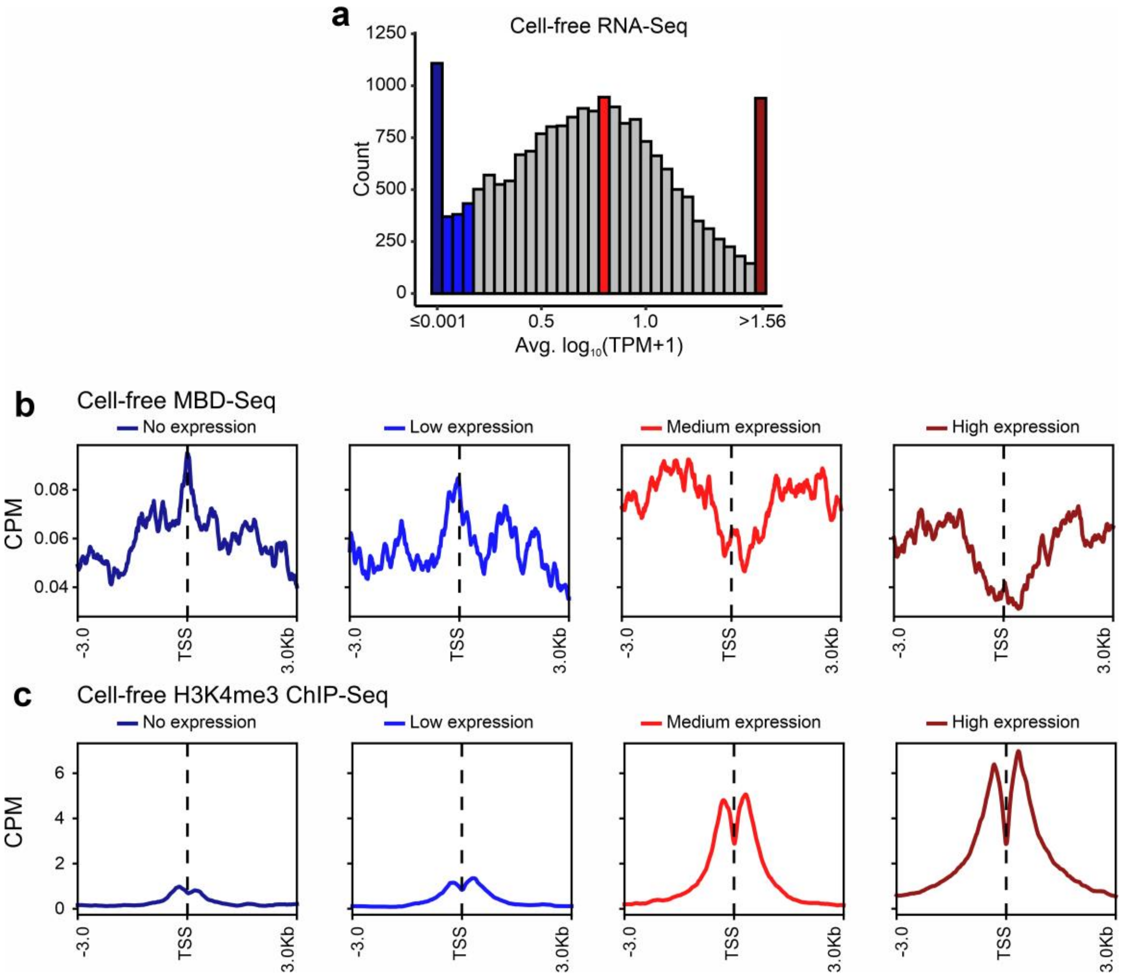


**Supplementary Figure S7 – Relationship of cell-free MBD-Seq and cell-free H3K4me3 ChIP-Seq with expression levels of genes.** (a) Histogram showing average expression levels of protein-coding genes in publicly available cell-free RNA sequencing data. For each category 938 genes were selected (5 % of all analyzed genes): no expression (dark blue), low expression (blue), medium expression (red), and high expression (dark red). (b) Average coverage profiles for cell-free MBD-Seq reads at selected transcription start sites. (c) Average coverage profiles for H3K4me3 ChIP-Seq reads at selected transcription start sites.

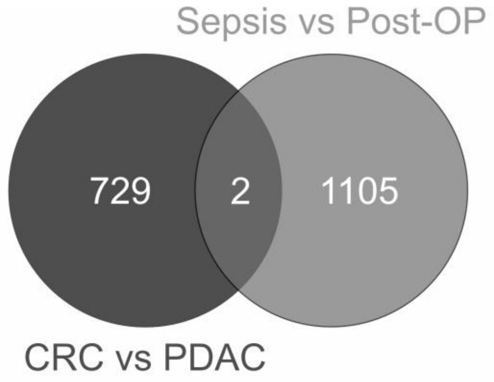


| Comparison | PDAC vsCRC | | Sepsis vsPost-OP | |
|---|---|---|---|---|
| High in | PDAC | CRC | Sepsis | Post-OP |
| # DERs | 543 | 188 | 663 | 444 |

**Supplementary Figure S8 – Comparison of loci with differential enrichment of footprint DNA.** The table shows the number of differentially enriched regions (DERs) derived from footprint DNA sequencing data for the comparisons of pancreatic ductal adenocarcinoma (PDAC) versus colorectal carcinoma (CRC) and post-operative controls (Post-OP) versus sepsis. The Venn diagram shows the overlap of the two sets of DERs.

**Supplementary Table S1 – Metadata for the generated patient sequencing datasets included in this study.** The sequenced datasets are available at the SRA (PRJNA1033613).

| Sequencing type | Sample | Condition | Age | Gender | Cancer stage |
|---|---|---|---|---|---|
| Liquid footprint | S01 | Healthy | 50 | male | - |
| Liquid footprint | S02 | Healthy | 50 | male | - |
| Liquid footprint | S03 | Healthy | 45 | female | - |
| Liquid footprint | S04 | Healthy | 45 | male | - |
| Regular cfDNA | S05 | Healthy | 47 | female | - |
| Regular cfDNA | S06 | Healthy | 61 | female | - |
| Regular cfDNA | S07 | Healthy | 50 | male | - |
| Regular cfDNA | S08 | Healthy | 52 | male | - |
| Liquid footprint | S09 | Post-OP | 20 | female | - |
| Liquid footprint | S10 | Post-OP | 38 | female | - |
| Liquid footprint | S11 | Post-OP | 29 | female | - |
| Liquid footprint | S12 | Post-OP | 56 | female | - |
| Liquid footprint | S13 | Sepsis | 50 | male | - |
| Liquid footprint | S14 | Sepsis | 65 | female | - |
| Liquid footprint | S15 | Sepsis | 74 | male | - |
| Liquid footprint | S16 | Sepsis | 71 | male | - |
| Liquid footprint | S17 | PDAC | 68 | male | III |
| Liquid footprint | S18 | PDAC | 75 | male | II |
| Liquid footprint | S19 | PDAC | 60 | male | I |
| Liquid footprint | S20 | PDAC | 63 | female | III |
| Liquid footprint | S21 | CRC | 84 | male | II |
| Liquid footprint | S22 | CRC | 65 | male | II |
| Liquid footprint | S23 | CRC | 72 | female | III |
| Liquid footprint | S24 | CRC | 86 | male | 0 (neoadjuvant treatment and full regression) |
| cfMBD-Seq | S25 | Healthy | 45 | female | - |
| Liquid footprint | S26 | Sepsis | 75 | male | - |
| Liquid footprint | S27 | Sepsis | 57 | male | - |
| Liquid footprint | S28 | Sepsis | 45 | male | - |
| Liquid footprint | S29 | Sepsis | 59 | male | - |
| cfMBD-Seq | S30 | Sepsis | 75 | male | - |
| cfMBD-Seq | S31 | Sepsis | 57 | male | - |
| cfMBD-Seq | S32 | Sepsis | 45 | male | - |
| cfMBD-Seq | S33 | Sepsis | 59 | male | - |
| RNA-Seq | S34 | Sepsis | 75 | male | - |
| RNA-Seq | S35 | Sepsis | 57 | male | - |
| RNA-Seq | S36 | Sepsis | 45 | male | - |
| RNA-Seq | S37 | Sepsis | 59 | male | - |

**Supplementary Table S2 – Metadata for the utilized public datasets in this study.** Cell-free H3K4me3 ChIP-Seq data are available from Zenodo: https://zenodo.org/record/4277001/files/Analysis.tgz?download=1. ATAC-Seq data from ENCODE were downloaded with the GRCh38 assembly and converted locally to the GRCh37 assembly with liftOver.

| Data type | Sample type | Source | Project ID | File ID |
|---|---|---|---|---|
| ATAC-Seq | Cell line (GM12878) | ENCODE | ENCSR095QNB | ENCFF646NWY |
| Cell-free RNA-Seq | Plasma from healthy individuals | SRA | PRJNA598835 | SRR10822588, SRR10822583, SRR10822579, SRR10822594, SRR10822591 |
| Cell-free H3K4me3 ChIP-Seq | Plasma from a healthy individual | Zenodo | - | H013.1 |
| DNase-Seq | Cell line (GM12878) | ENCODE | ENCSR000EMT | ENCFF783ZLL |
| DNase hypersensitive sites | Cell line (GM12878) | ENCODE | ENCSR000EMT | ENCFF273MVV |
| Double-stranded cell-free DNA | Plasma from a healthy individual | SRA | PRJNA291063 | SRR2130050 |
| Single-stranded cell-free DNA | Plasma from a healthy individual | SRA | PRJNA291063 | SRR2130051 |
